# The RES complex core subunits Snu17 and Bud13 buffer the spliceosome against intrinsically weak splicing signals in budding yeast

**DOI:** 10.64898/2026.10.19.760215

**Authors:** J. Angel Arce Quintor, Guillaume F. Chanfreau

## Abstract

Eukaryotic pre-mRNA splicing by the spliceosome requires a series of orchestrated rearrangements to promote formation of a catalytically active spliceosome. The Retention and Splicing (RES) complex, which contains Snu17p, Bud13p and Pml1p, assembles onto the B-complex and facilitates removal of the U4 snRNP while preventing premature catalytic activation of the spliceosome by Prp2p. To better understand the roles of each subunit of the RES complex *in vivo*, we analyzed splicing through RNA sequencing in *snu17*Δ, *bud13*Δ, and *pml1*Δ yeast mutants, with and without inactivation of nonsense-mediated decay. Inactivation of Snu17p and Bud13p led to a major decrease in splicing efficiency for many intron-containing genes, while inactivation of Pml1p affected only a very minor subset. Snu17p and Bud13p are particularly critical for promoting splicing of inefficiently spliced non-ribosomal protein genes with small introns, non-consensus 5′ splice sites or branchpoints, or without well-defined poly(U) tracts, suggesting a role for these elements in early intron recognition. The loss of these two core RES subunits therefore does not appear to cause splicing defects but rather exacerbate existing ones, such that transcripts with intrinsically high splicing efficiency remain unaffected by inactivation of RES. Cells lacking Pml1p also show growth phenotypes that are clearly distinct from those observed in *snu17*Δ and *bud13*Δ cells. These results show that Snu17p and Bud13p act as the core subcomplex of RES for efficient splicing of poorly spliced introns *in vivo*, while Pml1p may play accessory functions in linking the spliceosome to other processes.

## Introduction

The splicing of pre-mRNAs by the spliceosome is an essential and ubiquitous process in eukaryotic gene expression. Multicellular organisms diversify their proteome and regulate cell differentiation through alternative splicing switches that allow cells to produce proteins with different activities from the same pre-mRNA molecules and to regulate cell fate through these activities (Wilkinson et al, 2020; Merkin et al, 2012; Fiszbein & Kornblihtt 2017). For unicellular organisms such as *Saccharomyces cerevisiae*, RNA splicing is intimately linked with the proper maturation of highly expressed genes involved in meiosis, RNA transcription, RNA translation, and cytoskeletal dynamics, which equips cells with the ability to adapt to environmental cues by modulating the splicing of these transcripts (Wethekam & Moore 2023; Spingola et al. 1999; Parenteau et al. 2011).

In yeast, the earliest stage of intron recognition involves the U1 snRNP and the Msl5p-Mud2p heterodimer, which recognize and associate with the 5′ splice site (5′ss) and branchpoint (bp) sequences respectively during E complex formation (Wang et al. 2008; Wilkinson et al. 2020). The DEAD-box helicase Prp5p then triggers recruitment of the U2 snRNP by displacing Msl5p and Mud2p and allowing the U2 snRNA to base-pair with a stretch of about 14 nucleotides surrounding the bp adenosine area (Yan et al, 2026; Wilkinson et al, 2020), resulting in formation of the A complex. Recruitment of the U4/U6-U5 tri-snRNP follows and the B complex is formed when the helicases Prp28p and Brr2p sequentially displace the U1 and the U4 snRNPs, respectively. This rearrangement promotes the interactions between the 5′ss sequence and the U6 snRNA, as well as an extensive network of base-pairing interactions between the U6 and the U2 snRNAs that form the catalytic site of the spliceosome (Wilkinson et al. 2020). Throughout these transitions, the trimeric Retention and Splicing (RES) complex, which is comprised of Snu17p, Bud13p, and Pml1p is thought to maintain the structural integrity of the spliceosome and to prevent premature activation of the spliceosome (Bao et al. 2017). *In vitro*, premature binding of Prp2p in the absence of Snu17p causes spliceosome dissociation from intronic sequences, with only a small subset of structurally compromised spliceosomes able to complete the splicing reaction in a Prp2p-independent manner (Bao et al. 2017).

NMR studies have shown that the RES complex is anchored directly to the pre-mRNA by the RNA Recognition Motif (RRM) of Snu17p, which tightly associates with intronic sequences in the presence of Bud13p and, potentially, Pml1p (Wysoczanski et al. 2015). Cryo-EM structures of the Bact spliceosome revealed that the palm of Snu17p binds about 16 nucleotides downstream of the branchpoint adenosine, covering a stretch of 6 nucleotides (Fig.1A) immediately downstream of Hsh155p (Yan et al. 2026). Pml1p is anchored to the spliceosome through a weak interaction with Snu17p (Yan et al. 2026), which might be stabilized by Prp45p although this interaction has not been biochemically tested. Prp45p loops entirely around the RES complex through a curl that begins with a direct interaction with a region on the N-terminal end of Pml1p and follows towards the C-terminal end of Prp45p in a lasso that fully wraps around Bud13p and then Snu17p (Yan et al. 2026; Fig. 1B). Prp45p also sits over the FHA domain of Pml1p and “clamps” it in place, potentially preventing it from dissociating from the rest of the RES complex (Fig. 1C).

**Figure 1.**
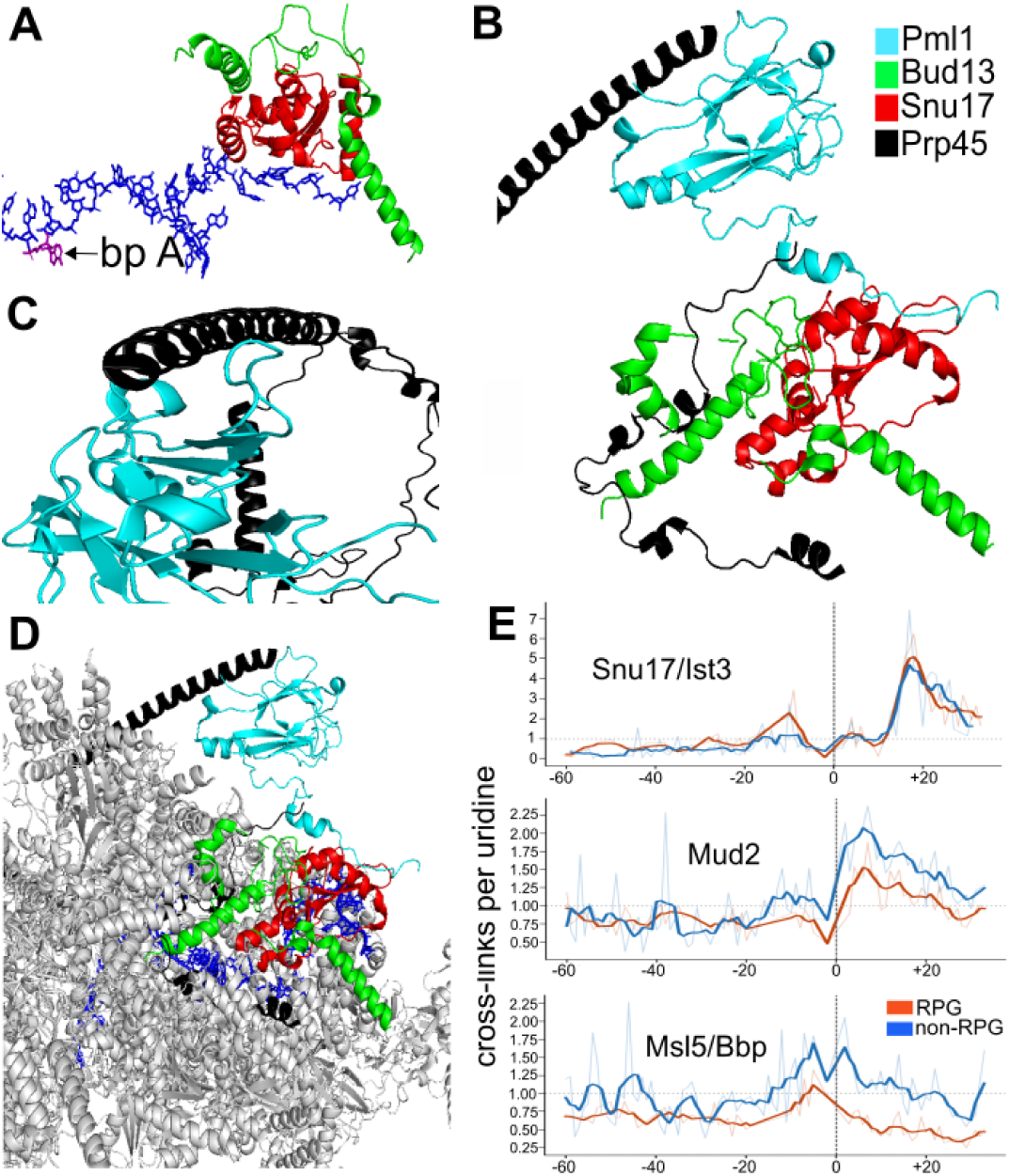
Structural and cross-linking data show that Snu17p binds pre-mRNA ∼17 nucleotides downstream of the bp. The proteins Prp45p, Snu17p, Bud13p and Pml1p are color-coded in black, red, green, and cyan, respectively. **A.** Snu17p-Bud13p heterodimer binding pre-mRNA downstream of the bp. The bp A is highlighted in purple. **B.** The main *α*-helix of Prp45p stabilizes the Snu17p-Pml1p interaction by clamping the FHA domain of Pml1p to the body of the spliceosome. Prp45p wraps around the main body of the RES complex and interacts mostly with Bud13p. **C.** Interaction interface between Prp45p and Pml1p. **D.** Global position of the RES complex in the spliceosome. Snu17p and Bud13p sit in the main body of the spliceosome while the FHA domain of Pml1p sits outside. **E.** Uridine-normalized PAR-CLIP cross-link profiles of Snu17p/Ist3p, Mud2, and Msl5p (top to bottom) plotted relative to the branchpoint adenosine (dashed line = 0). Cross-links per uridine are scaled to its own intron-wide mean (dotted line = 1); averaged separately between RPGs (red, n = 89) and all others (blue, n = 149).

Pml1p contains a Forkhead-associated (FHA) domain (Trowitzsch et al. 2009). It sits outside the main body of the spliceosome (Fig. 1C and 1D) and likely recognizes phosphorylated residues on proteins at the vicinity of the spliceosome (Mahajanet al. 2008; Trowitzsch et al. 2009). This domain therefore could mediate communication between spliceosome components and other proteins involved in nuclear RNA maturation and transport. This is corroborated by the observation that deletion of *PML1* does not affect the splicing of an in-frame pre-mRNA reporter in yeast but causes the leakage of the unspliced transcript into the cytoplasm (Dziembowski et al. 2004). It is also supported by the recent discovery that the FHA domain of Pml1p mediates phosphorylation of the core spliceosome component SF3b in humans by the cyclin-dependent kinase 11 (CDK11), a reaction necessary for the Bact to B* transition in mammals (Gajdušková et al. 2026; Trowitzsch et al. 2009).

Snu17p, Bud13p and Pml1p are evolutionarily conserved from yeast to humans (Dziembowski et al. 2004). In zebrafish, the RES complex is enriched in the central nervous system during early development with zebrafish loss-of-function mutants in s*nu17*, *bud13* and *pml1* showing severe developmental defects of the central nervous system and reduced splicing efficiency (SE) of most intron-containing genes (Fernandez et al, 2018). Short introns with high GC content are the most affected in the absence of functional Snu17p and Bud13p and Pml1p, indicating that the RES complex is important, at least in this organism, for the removal of introns excised through an intron-definition mechanism (Fernandez et al. 2013). In *Arabidopsis thaliana*, the protein products of *GDS1*, *AtBUD13* and *DDL1*, the orthologues of *SNU17*, *BUD13* and *PML1*, respectively, have been shown to associate with one another in a two-hybrid system and contribute as a complex to the efficient splicing of a small subset of transcripts that regulate embryo development and root growth (Xiong et al. 2022). RNA-seq analysis of homozygous *atbud13* mutants shows decreased expression of at least 52 transcripts harboring small intronic sequences (<100 nucleotides), likely due to the generation of unspliced transcripts with premature termination codons (Xiong et al. 2019).

Whether or not each subunit of RES plays a distinct role in splicing *in vivo* is currently not well understood. In *Saccharomyces cerevisiae*, Snu17p and Bud13p, but not Pml1p, have been implicated in the Mer1p-dependent splicing of meiotic transcripts with weak 5′ss and bp sequences (Spingola et al. 2004; Scherrer et al. 2006). Splicing of *MED20* and *TAN1* is dependent on the RES complex, with both transcripts showing almost complete abrogation of splicing in *snu17*Δ and *bud13*Δ strains, but not *pml1*Δ (Zhou et al. 2013; Zhou & Johansson 2017). Both *MED20* and *TAN1* harbor small intronic sequences (< 75 nts) and the branchpoint sequence of *MED20* deviates from the consensus sequence. Because Med20p is important for transcriptional activation and Tan1p is essential for tRNA modifications, the growth defect observed in *snu17*Δ and *bud13*Δ strains at 37 ℃ has been attributed primarily to general defects in the transcription and tRNA processing machineries (Zhou et al, 2013; Zhou & Johansson 2017). Nonetheless, pronounced decreases in SE for these two transcripts has been observed in a wide range of other splicing mutant datasets such as *prp18*Δ, *luc7*Δ, *prp45*Δ, *mud1*Δ, *isy1*Δ, *prp17*Δ and others, which indicates that these two transcripts are poorly spliced in many other splicing factor mutants and not just in RES mutants (Abrhámová et al. 2024; Kawashima et al. 2014; Carrocci et al. 2024).

To understand the role of each subunit of RES in splicing *in vivo*, we first clarified the positioning of Snu17p on intronic sequences *in vivo* and analyzed splicing genome-wide through RNA sequencing of strains lacking each subunit of the complex. Reanalysis of published PAR-CLIP data positioned Snu17p binding to RNA approximately 17 nucleotides downstream of the branch-point adenosine, consistent with recent cryo-EM structures of the Bact spliceosome. RNA sequencing of mutant strains showed that while Pml1p is largely dispensable for splicing, loss of Snu17p or Bud13p lowers splicing efficiency genome-wide, with strikingly similar effects. Pml1p deletion produces growth phenotypes and a strong synthetic interaction with inactivation of nonsense mediated decay that are distinct from those of the other two mutants. Our results show that Snu17p and Bud13p act as the functional core of RES, while Pml1 may play accessory roles outside splicing.

## Results

### Snu17p binds introns 17 nucleotides downstream of the branch-point adenosine *in vivo*

The RNA recognition motif (RRM) of Snu17p is essential in tethering the RES complex to the pre-mRNA prior to the first transesterification reaction (Yan et al. 2008). Cryo-EM structures show Snu17p directly associating with the pre-mRNA exactly 16 nucleotides downstream of the bp A, which accords with the observed enrichment of U2 components in Snu17p immune-precipitation experiments (Fig. 1A; Yan et al. 2008; Gottschalk et al. 2001). On the other hand, PAR-CLIP studies of RNA processing factors using 4-thiouridine incorporation have mapped the RRM of Snu17p to an area that peaks 27 nucleotides downstream from the branchpoint and immediately downstream of the apparent Mud2p binding site (Baejen et al. 2014). Although the authors concluded that the binding site of Snu17p is restricted by the location of Mud2p on the pre-mRNA, this positioning by Mud2p is highly unlikely, as Mud2p is displaced from the spliceosome by Prp5p before A complex formation, while Snu17p only associates with the pre-mRNA during B-complex formation, well after the joining of the U4/U6-U5 tri-snRNP (Wang et al. 2008; Bao et al. 2017; Winkinson et al. 2020). Counting polyU content generates a peak exactly 27 nucleotides downstream of the bp A (Beajen et al. 2014), suggesting that the apparent peak of Snu17p at this position was due to polyU content downstream of the bp sequence. We therefore re-analyzed cross-linking data of Msl5p, Mud2p and Snu17p (named Ist3p in that study), with Msl5p acting as an internal control whose signal should peak at Us within the bp sequence. For each annotated intron, we computed the mean number of T-C transitions per uridine across the entire intron length, and at each position containing a uridine, we divided the transition count by the introns’ mean. At each position relative to the bp adenosine, normalized values were averaged across all introns containing a uridine at that position, including those that showed no transitions. Introns that did not contain a uridine at the specified position were excluded from the analysis. Finally, positions supported by less than 30 introns were not included in the metagene plots. Using this uridine-based normalization, we found that the Msl5p crosslinks peaked upstream of the bp A at positions -5 and -2, owing to the uridines in the bp sequence (<u>U</u>ac<u>U</u>aac). Non ribosomal protein genes (Non-RPGs) showed an additional peak at position +1 relative to the bp A, which is caused by non-consensus UacUaaU bp sequences found in several genes such as *HRB1*, *SCS22*, *YKR005C* and *GMC2* (Fig. 1E). The crosslink positions of Mud2p and Snu17p/Ist3p shifted towards the 5′ end of the intron by 9 and 10 nucleotides, respectively, relative to the positions reported by Beajen *et al*. This analysis showed that Snu17p crosslinks peaks exactly 17 nucleotides downstream of the bp A (Fig. 1E), a position which is in perfect agreement with the position of Snu17p established by the published structures of the Bact spliceosome (Rauhut et al. 2016; Yan et al. 2016). Thus, our analysis supports the model where Snu17p/Ist3p associates with the pre-mRNA during B complex formation and suggests that its position is determined primarily by its association with Hsh155p.

### Inactivation of Snu17p or Bud13p, but not Pml1p, causes major reductions in splicing efficiency in *Saccharomyces cerevisiae*

We next sought to analyze the effect inactivation of individual RES components has on splicing efficiency *in vivo*. Determining the splicing efficiency of intron-containing genes (ICGs) genome-wide in splicing mutants is complicated by the activity of the nonsense-mediated decay (NMD) pathway, which degrades unspliced pre-mRNAs retaining premature termination codons (PTCs) (He et al. 1993; Sayani et al. 2008). To accurately quantify SEs across the yeast transcriptome in the absence of each RES subunit, we deleted *UPF1* in wt, *snu17*Δ, *bud13*Δ and *pml1*Δ cells and performed total RNA sequencing in both NMD-competent and NMD-deficient backgrounds. During library preparations, one RNA sample from a *pml1*Δ replicate was inadvertently replaced with one *pml1*Δ*upf1*Δ RNA, resulting in two *pml1*Δ replicates and four *pml1*Δ*upf1*Δ replicates. Reads were mapped with BBMAP and we obtained an average of 48M uniquely mapped reads per individual replicate. We then used SPLICEq (Costa et al. 2020) on our aligned BAM files and calculated the SE of annotated ICGs with enough read coverage in the strains deficient for NMD. PCA analysis of the SE showed high agreement between sample replicates, including the fourth *pml1*Δ*upf1*Δ replicate which clustered close to the other 3 replicates of the same strain (Fig. 2C). All subsequent analyses were performed with four *pml1*Δ*upf1*Δ replicates and two *pml1*Δ replicates.

**Figure 2.**
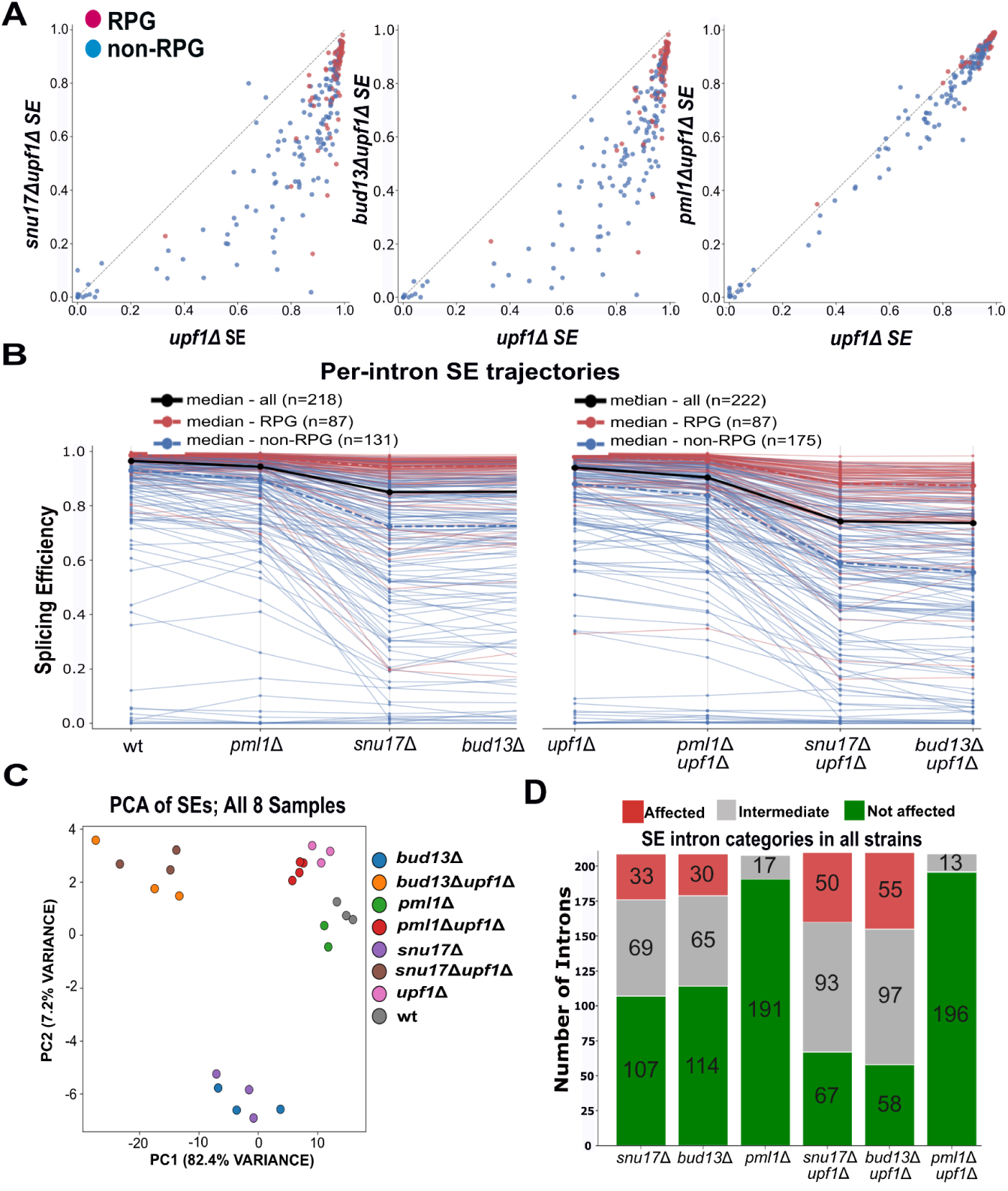
Bioinformatics analysis of splicing efficiencies (SE) in wt, *upf1*Δ, RES single mutants and RES-*upf1*Δ double mutant strains. **A.** Scatter plots comparing the SEs of *snu17*Δ*upf1*Δ, *bud13*Δ*upf1*Δ and *pml1*Δ*upf1*Δ against *upf1*Δ. Individual points represent annotated junctions detected with SPLICEq. RPGs are color-coded as red while non-RPGs are blue. Points below the diagonal are splice junctions whose SE is lower upon deletion of a RES complex component. **B.** Parallel trajectory plots of the calculated SE of individual junctions. The median SEs of RPGs and non-RPGs are shown in dotted red and blue, respectively. Global median SE is shown in solid black. **C.** PCA analysis of calculated SEs in all 8 strains. **D.** Introns categorization based on decreases in SE upon RES-complex component deletion. Introns with a ΔSE ≤ 0.1 are categorized as unaffected. Introns with a ΔSE ≥ 0.3 are categorized as affected. Introns with a ΔSE in between 0.1 and 0.3 are categorized as intermediate.

Deletion of *SNU17* and *BUD13* led to significant SE decreases genome-wide, with pre-mRNAs of non-RPGs being the most affected (Fig. 2A). By contrast, deletion of *PML1* did not result in any major changes in SE (Fig. 2A), a result consistent with the peripheral position of Pml1p outside of the main body of the Bact spliceosome and away from the pre-mRNA (Fig. 1D; Yan et al. 2016). To compare the effect of each RES component on individual ICG SE, we plotted the calculated SEs of all strains on a parallel trajectory plot (Fig 2B). Pml1p inactivation showed only a mild effect on the median SE of all calculated introns (1.1% for RPGs and 3.7% for non-RPGs). By contrast, inactivation of Snu17p and Bud13p led to a decrease in the median SE of RPGs of 9.5% and 10.1%, respectively. Non-RPGs were significantly more affected with 30.2% and 32.9% decrease upon Snu17p or Bud13p inactivation in the NMD-deficient background. RPGs were overall more resistant to defects in splicing upon deletion of either one of these three proteins, an effect that has been observed in other splicing factor mutants (Pleiss et al. 2007; Roy et al. 2023). Non-RPG introns were slightly more affected by the absence of Bud13p than Snu17p (Fig. 2B). The same trends were observed for the single mutants compared to wild-type, although less pronounced due to the activity of NMD masking unspliced the accumulation of unspliced RNAs (Supplementary Fig. 1B).

We then classified ICGs according to their SE changes (ΔSE) upon each RES subunit inactivation in both wt and *upf1*Δ backgrounds. Unaffected introns were defined as showing less than a 10% decrease in SE, intermediates having a ΔSE between 10 and 30%, and strongly affected as showing a ΔSE higher than 30% upon deletion of a given RES subunit. Nearly half of all ICGs appeared as unaffected in *snu17*Δ and *bud13*Δ in a wt background. This number decreased sharply upon NMD inactivation from 107 to 67 in *snu17*Δ*upf1*Δ and from 114 to 58 in *bud13*Δ*upf1*Δ (Fig. 2D), consistent with NMD masking a substantial fraction of splicing defects in the single RES mutants. By contrast, the number of unaffected introns in *pml1*Δ remained essentially unchanged whether or not NMD was active (191 in *pml1*Δ to 196 in *pml1*Δ*upf1*Δ), indicating that *PML1* inactivation generates very few significant splicing defects. These results establish that the deletion of *SNU17* and *BUD13*, but not *PML1*, reduce the SEs of many ICGs and define for each subunit the specific subset of introns most sensitive to their loss (Fig. 2B).

### PolyU tracts buffer SE decrease in mutants of the core RES complex

To understand how splice signals affect changes in SE upon RES inactivation, we generated sequence logos from the 5′ss, bp, and 3′ss sequences in the different classes of ICGs defined above as affected, moderately affected, or unaffected by inactivation of each component of RES (Fig. 2D). In the *snu17*Δ*upf1*Δ and *bud13*Δ*upf1*Δ mutants, a well-defined polyU tract upstream of the 3′ss differentiates unaffected introns from the other two categories while no difference in 5′ss sequence was observed between ICG classes (Fig. 3). The bp sequence of affected introns in *snu17*Δ*upf1*Δ and *bud13*Δ*upf1*Δ showed less constraint for a U at position -5 relative to the bp adenosine compared to unaffected and intermediate categories. Unaffected introns also showed an enrichment of U at position -5 of the bp compared to the overall ICG population (Fig. 3). This depletion of uridine at position -5 in the bp sequence among introns that are affected in *snu17*Δ*upf1*Δ and *bud13*Δ*upf1*Δ is consistent with the observation that a small subset of meiotic transcripts harboring non-consensus bp sequences are among the most affected upon deletion of *SNU17* and *BUD13* (Spingola et al. 2004; Scherrer & Spingola 2006). Introns that are strongly or moderately affected by deletion of *SNU17* and *BUD13* also exhibit weaker polyU tracts compared to unaffected introns. This result supports a model where a well-defined polyU tract between the bp and the 3′ss buffers introns against loss of either Snu17p and Bud13p—the two RES subunits that contact this region in the pre-mRNA (Wysoczanski et al. 2015; Yan et al. 2026). We could not test this directly for Pml1p as no introns showed decreases in SE greater than 30% after its deletion. The polyU tracts detected among *pml1*Δ-unaffected introns likely reflect sequence bias present across the bulk of ICGs rather than a Pml1p-specific effect.

**Figure 3.**
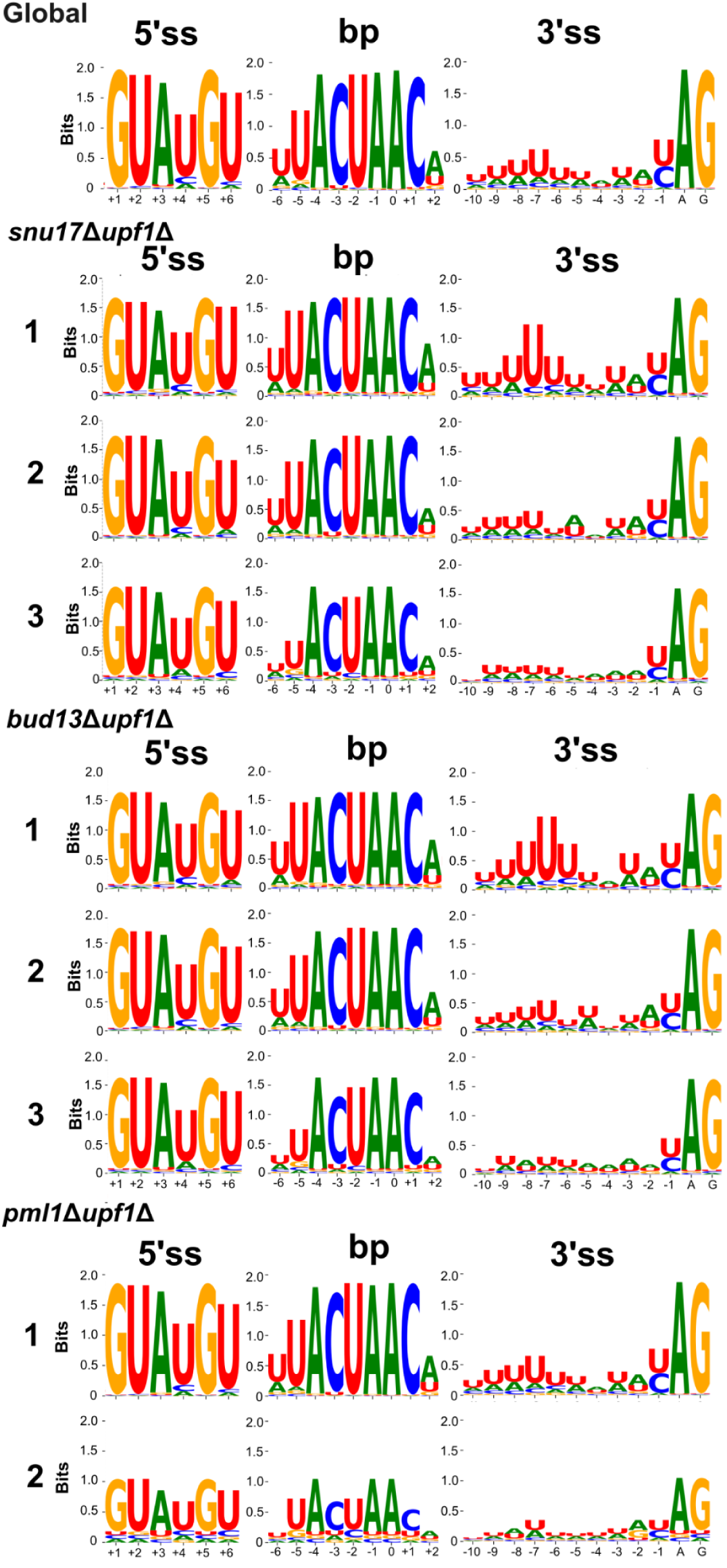
Splice-site sequence features of introns grouped by sensitivity of SE to the inactivation of each RES subunit. Sequence logos of the 5′ splice site, branchpoint and 3′ splice site are shown. The top panel shows logos for all annotated *S. cerevisiae* introns as a reference. The remaining panels show logos for introns classified as unaffected (**1**), intermediately affected (**2**), and affected (**3**) in *snu17*Δ*upf1*Δ, *bud13Δupf1Δ*, and *pml1Δupf1Δ*. Introns were classified by ΔSE values as described in Figure 1. No logos are shown for affected introns (cat 3) in *pml1Δupf1Δ* because no introns met the criteria for that category. Letter height is information content in bits.

### Random Forest Permutation analysis identifies branch-point deviations and RPG status as major determinants of splicing efficiency in RES mutants

To further identify intronic or genomic features that predict splicing efficiency (SE) in the absence of each RES subunit, we performed a Random Forest permutation-importance analysis (Gregorutti et al. 2016).

For every ICG with a mapped transcriptional start site (TSS) (McMillan et al. 2019), we defined nine intronic features: distances from TSS-to-5′ss, TSS-to-bp, TSS-to-3′ss, and bp-to-3′ss; deviations of 5′ss or bp sequences from the consensus; intron length; polyU content between the bp and the 3′ss; and the longest U-stretch in that same interval. We added RPG status as a tenth feature, since RPGs have systematically longer introns, shorter TSS-to-5′ss distances, and longer TSS-to-bp and TSS-to-3′ss distances than non-RPGs (Supplemental Figure 1A). We also included three splicing kinetics parameters previously measured *in vivo* at 20°C in *S.cerevisiae*, the half-lives of the first-step, second-step, and total splicing reaction (Gildea et al. 2022). This generated a total of 13 predictors. Each feature was permuted 50 times and ranked by its mean decrease in model R². To remove the confounding effect of NMD, our discussion is focused exclusively on the results obtained in the NMD-deficient background, although the analysis done in wt background strains corroborates the observed results in the *upf1*Δ background (Supplemental Figure 1B). The results for each strain are presented in Fig.4 A-D.

**Figure 4.**
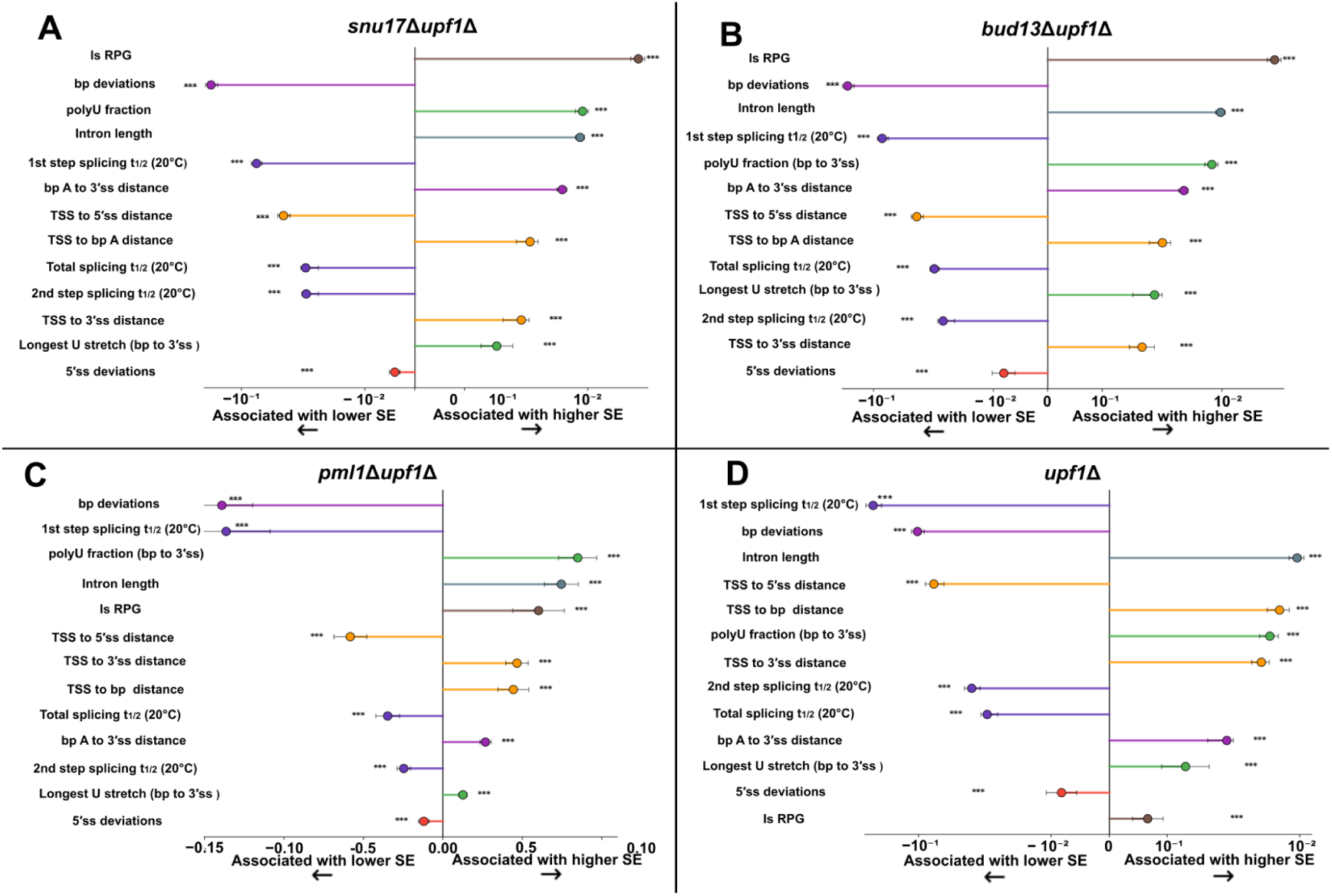
Random Forest permutation importance for SE in *upf1*Δ background strains. Each feature was independently shuffled a total of 50 times. The length of the lollipop stem shows the mean decrease in model R². Defined features associated with low SE values are shown on the left of each plot while those associated with high SE values are shown on the right of each plot. Statistical significance of each feature’s importance was assessed with a one-sided Wilcoxon signed-rank test across permutation repeats (H₀: importance ≤ 0), with raw p-values corrected for multiple comparisons within each model by the Benjamini-Hochberg procedure (*** FDR < 0.001, ** FDR < 0.01, * FDR < 0.05). Sample sizes, R², and model p-value for each genotype were: *upf1*Δ, n = [191], R² = [0.132], p = [0.01]; *snu17*Δ*upf1*Δ, n = [191], R² = [0.370], p = [0.01]; *bud13*Δ*upf1*Δ, n = [191], R² = [0.408], p = [0.01]; *pml1*Δ*upf1*Δ, n = [190], R² = [0.238], p = [0.01]. Shown are features ranked by permutation importance for each mutant. **A.** *snu17*Δ*upf1*Δ **B.** *bud13*Δ*upf1*Δ **C.** *pml1*Δ*upf1*Δ and **D.** *upf1*Δ.

In *upf1*Δ (pseudo-WT in this context), the first-step splicing half-life was the strongest predictor of SE, with long half-lives associated with low SE (Fig. 4D). This low SE reflects accumulation of unspliced precursors in the absence of functional NMD for unspliced transcripts. Deviations from the consensus bp sequence and intron length ranked second and third, with bp deviation lowering SE, and longer introns splicing more efficiently. TSS-to-5′ss distance ranked fourth with longer distances reducing SE. This observation is consistent with recent findings that for a given gene, transcripts with long 5′-exons show lower SE than shorter ones (Barr et al. 2025). TSS-to-bp distance, polyU fraction, and TSS-to-3′ss distance followed as the next-best predictors, all three positively associated with high SE. Second-step and total splicing half-life were also negatively associated with SE, though both ranked below first-step half-life. RPG status ranked last of the 13 predictors in *upf1*Δ.

By contrast, RPG status was the single best predictor of high SE in *snu17*Δ*upf1*Δ and *bud13*Δ*upf1*Δ (Fig. 4A, B). Most RPGs remained efficiently spliced in the absence of Snu17p or Bud13p, indicating that neither subunit is critical for efficient splicing; a small subset of RPGs is nonetheless affected by their absence (Supplemental Figure 2A). Interestingly, deviations from the consensus 5′ss was a poor predictor, which might be linked to RPG status. By splitting intron-containing RPGs into those harboring a consensus 5′ss (GUAUGU) and those lacking a consensus 5′ss, we observed no differences in the median SE in any of our conditions (Supplemental Figure 2B). Non-RPGs showed the opposite pattern, with non-consensus 5′ss sequences being associated with lower SE compared to consensus ones (Supplemental Figure 2C). Bp deviation ranked second in both mutants (Fig. 4A, B) and associated with low SE. PolyU fraction and intron length ranked third and fourth (order reversed between the two strains), both raising SE, while TSS-to-5′ss distance and 5′ss deviation were associated with lower SE in both strains, ranking lower overall. The 13 predictors had similar ordering and impact sizes for the *snu17*Δ*upf1*Δ and *bud13*Δ*upf1*Δ mutants (Fig. 4A & B), consistent with the idea of Snu17p and Bud13p acting as core components of a single functional unit in the spliceosome. Splicing half-lives showed the same effect as in the *upf1*Δ strain, with a long first-step half-life again being the strongest predictor of low SE.

In *pml1*Δ*upf1*Δ, the top two predictors were deviation from the bp sequence and first-step half-life—both associated with lower SE. PolyU fraction, intron length, and RPG status were associated with higher SE (Fig. 4C). These results show that deviation from the consensus bp sequence ranks highly across all three RES/*upf1*Δ strains but that RPG status ranks lower in *pml1*Δ*upf1*Δ than in *snu17*Δ*upf1*Δ and *bud13Δupf1*Δ. This confirms that deletion of *SNU17* and *BUD13* widens the difference in SE between RPGs and non-RPGs more dramatically than deletion of *PML1,* and shows that predictors such as bp deviations, polyU fraction, intron length and 1st step splicing kinetics rank highly in all three strains.

### Small introns and introns inefficiently spliced at background levels show the greatest decrease in SE upon Snu17p or Bud13p inactivation

We expanded the Random Forest Permutation analysis by calculating the change in SE (ΔSE) of individual introns upon RES inactivation as the difference between the SE in the RES-NMD double KO strains and the SE in the *upf1*Δ control strain. This second Random Forest Permutation was performed using the same predictors described above to understand what genomic features contribute to the largest decreases in SE in the absence of each subunit of the RES complex. We included background SE (SE in *upf1*Δ) as a predictor to accommodate for the observation that small introns and non-RPGs with non-consensus 5′ss and bp signals are already poorly spliced in *upf1*Δ.

Background SE ranked as the top predictor of ΔSE in every RES-NMD double KO strains (Fig. 5A, B & C), showing that introns with intrinsically low background SE are more likely to show high decrease in SE upon RES inactivation. This result suggests that loss of Snu17p or Bud13p exacerbates defects in splicing instead of causing them. This is illustrated by *TAN1* and *MED20*: these transcripts already exhibit a mild splicing defect in the background strain and splicing is completely abrogated in *snu17*Δ*upf1*Δ and *bud13*Δ*upf1*Δ. By contrast, *ACT1* and *TUB2* show no defects in splicing in control strains and do not exhibit high levels of unspliced transcripts in the *snu17*Δ*upf1*Δ and *bud13*Δ*upf1*Δ mutants (Fig 5D).

**Figure 5.**
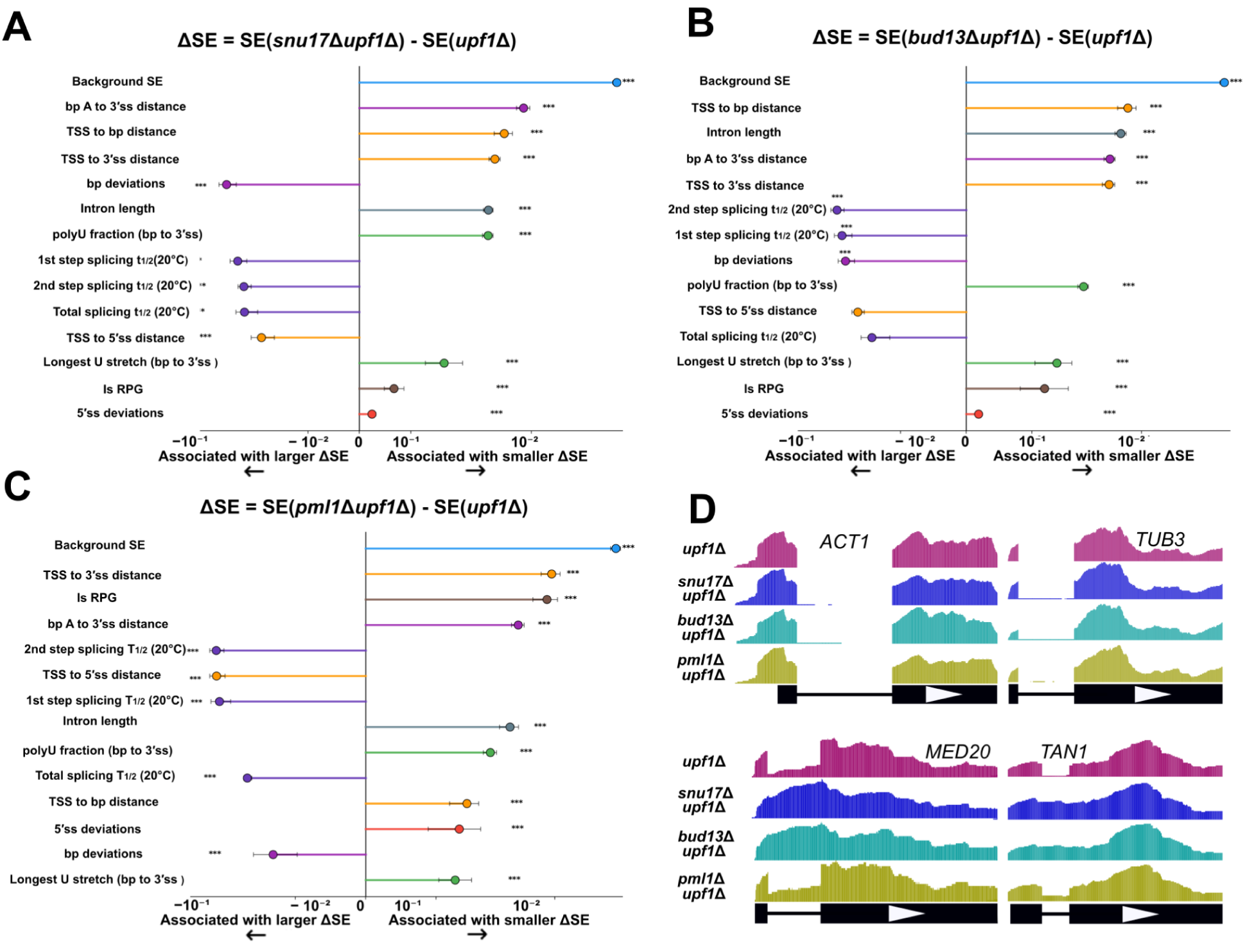
Random Forest permutation analysis of ΔSE in *upf1*Δ background strains. The analysis was performed as described in Figure 3 with the addition of background SE (SE in *upf1*Δ). The decrease in SE (ΔSE) for each annotated junction was calculated for all three RES-NMD double KO strains as **A.** SE*snu17*Δ*upf1*Δ – SE*upf1*Δ, **B.** SE*bud13*Δ*upf1*Δ – SE*upf1*Δ and **C.** SE*pml1*Δ*upf1*Δ – SE*upf1*Δ. Sample sizes, R², and model p-value for each genotype were: *upf1*Δ, n = [191], R² = [0.132], p = [0.01]; *snu17*Δ*upf1*Δ, n = [191], R² = [0.151], p = [0.01]; *bud13*Δ*upf1*Δ, n = [191], R² = [0.243], p = [0.01]; *pml1*Δ*upf1*Δ, n = [190], R² = [0.147], p = [0.01]. **D.** The introns *MED20* and *TAN1* have large decreases in SE upon deletion of RES complex components while the *ACT1* and *TUB2* (both non-RPGs), have negligible ΔSE.

In *snu17*Δ*upf1*Δ, bp-to-3′ss distance ranked as the second-best predictor of ΔSE, followed by TSS-to-bp and TSS-to-3′ss distance; longer distances in all three features were associated with a smaller ΔSE. Deviation from the consensus bp sequence ranked fifth, associated with a larger decrease in SE, followed by intron length and polyU fraction between the bp and 3′ss. TSS-to-5′ss distance ranked lower; longer distances were associated with a larger ΔSE. In *bud13*Δ*upf1*Δ, TSS-to-bp distance ranked as the second-best predictor of ΔSE, followed by intron length and bp-to-3′ss distance; bp deviation ranked lower here than in *snu17*Δ*upf1*Δ. The results indicate that intron length carries more importance in *bud13*Δ*upf1*Δ than in *snu17*Δ*upf1*Δ (0.059 vs. 0.036). Plotting the median ΔSEs of all introns with less than 100 nucleotides in length in all RES-NMD KOs reveals that small introns are more susceptible to deletion of *BUD13* than to deletion of *SNU17* (Supplemental Figure 3A). This difference is small, however, and not statistically significant. Splicing kinetics half-lives ranked in the middle of both strains’ predictor lists, with large values for all three half-lives negatively associated with ΔSE, as expected. Lastly, RPG status and deviations from the 5′ss ranked the lowest in both strains.

By contrast, RPG status ranked highly as a predictor of small ΔSE change in *pml1*Δ*upf1*Δ, just below background SE and TSS-to-3′ss distance. This result shows that RPGs and introns with long TSS-to-3′ss and bp-to-3′ss spacing were protected from the mild effect of *PML1* deletion. Splicing half-lives and TSS-to-5′ss distance occupied the middle ranks, largely mirroring their roles in the other two strains. Bp deviation and intron length, which ranked prominently in *snu17*Δ*upf1*Δ and *bud13*Δ*upf1*Δ, fell to the bottom third of predictors here, once again underscoring the differences between *pml1*Δ*upf1*Δ and *snu17*Δ*upf1*Δ and *bud13*Δ*upf1*Δ.

Together, our results indicate that a subset of small, non-RPG ICGs with short bp-to-3′ss distances, non-canonical bp sequences, and slow intrinsic splicing kinetics are already poorly spliced at background levels. These transcripts are the most susceptible to inactivation of Snu17p or Bud13p, but much less to Pml1p. Loss of the core two RES complex subunits therefore does not appear to intrinsically cause splicing defects but rather to exacerbate existing ones. This means that transcripts very efficiently spliced at background levels are impervious to deletion of either one of these three proteins.

### *SNU17* and *BUD13* deletions downregulate overall transcript levels of ICGs and RPGs

We next analyzed the impact of RES subunits inactivation on gene expression, focusing on the effect on ICGs. Using DESeq2 (padj < 0.05; Fisher’s exact test), we found that inactivation of Snu17p or Bud13p had a major impact on the expression levels of ICGs, while Pml1p inactivation had a much more modest effect (Supplemental Fig. 3B; Supplemental table 2). In *snu17*Δ, 19.5% of all ICGs were significantly downregulated, 3.8% increased in expression, and 76.7% remained unchanged. In *bud13*Δ, 18.8% of ICGs decreased in expression, 3.8% increased, and 77.4% showed no significant change. By contrast 92.1% of ICGs showed no change in expression in *pml1*Δ (Supplemental Fig. 3B), in agreement with the idea that Snu17p and Bud13p act as the core components of RES. The few ICGs that increased in transcript level in *snu17*Δ and *bud13*Δ correspond to genes hosting intronic snoRNAs, such as *NOG2*. snoRNA-host ICGs were previously shown to accumulate hybrid mRNA-snoRNA transcripts (hmsnoRNAs) upon splicing inactivation (Liu et al. 2022), and the apparent increase in overall transcript levels of these genes in *snu17*Δ and *bud13*Δ likely reflects a strong accumulation of hmsnoRNAs rather than an actual increase in spliced mRNA (Supplemental Fig. 3C).

Surprisingly, many of the ICGs downregulated in the absence of Snu17p or Bud13p are RPGs which accumulate as fully spliced transcripts even in the absence of these subunits (e.g. *RPL7A*, Supplemental Fig. 3D). A decrease in transcript levels in the *snu17*Δ and *bud13*Δ strains is therefore inconsistent with their apparent efficient splicing, raising the question of the mechanism involved in this decrease in expression. We analyzed this further by performing a global correlation analysis between SE and transcript expression (restricted to the NMD-deficient backgrounds, since SE cannot be accurately estimated when unspliced transcripts are subject to NMD), and found that large decreases in SE correlate negatively with the expression of individual transcripts, an effect that was statistically significant in *snu17*Δ*upf1*Δ, *bud13*Δ*upf1*Δ, and *pml1*Δ*upf1*Δ (Fig. 6A–C, right panels). Surprisingly, intron-containing RPGs’ abundance decreased more dramatically than non-RPGs’ upon deletion of either RES subunit; an effect that’s at odds with their observed low decreases in SE. This led to the hypothesis that deletion of *SNU17*, *BUD13* or *PML1* triggers a transcriptional feedback loop that reduces global RPG expression. To test this hypothesis, we split the full set of annotated RPGs in *S. cerevisiae* (n = 138) into intron-containing (102) and intronless (36) genes and compared their expression in each RES single mutant (Fig. 6A–C, left panels). Intronless RPGs were downregulated to levels statistically indistinguishable from those of intron-containing RPGs in all three mutants, including *pml1*Δ. This indicates that a transcriptional feedback mechanism is triggered by deletion of any of the three RES genes, reducing RPG expression independently of splicing status. By contrast, intron-containing non-RPGs showed significant decrease in transcript levels in the Snu17p or Bud13p mutants, but not in the Pml1p mutant, which suggests that these decreases in expression are directly related to their splicing defects.

**Figure 6.**
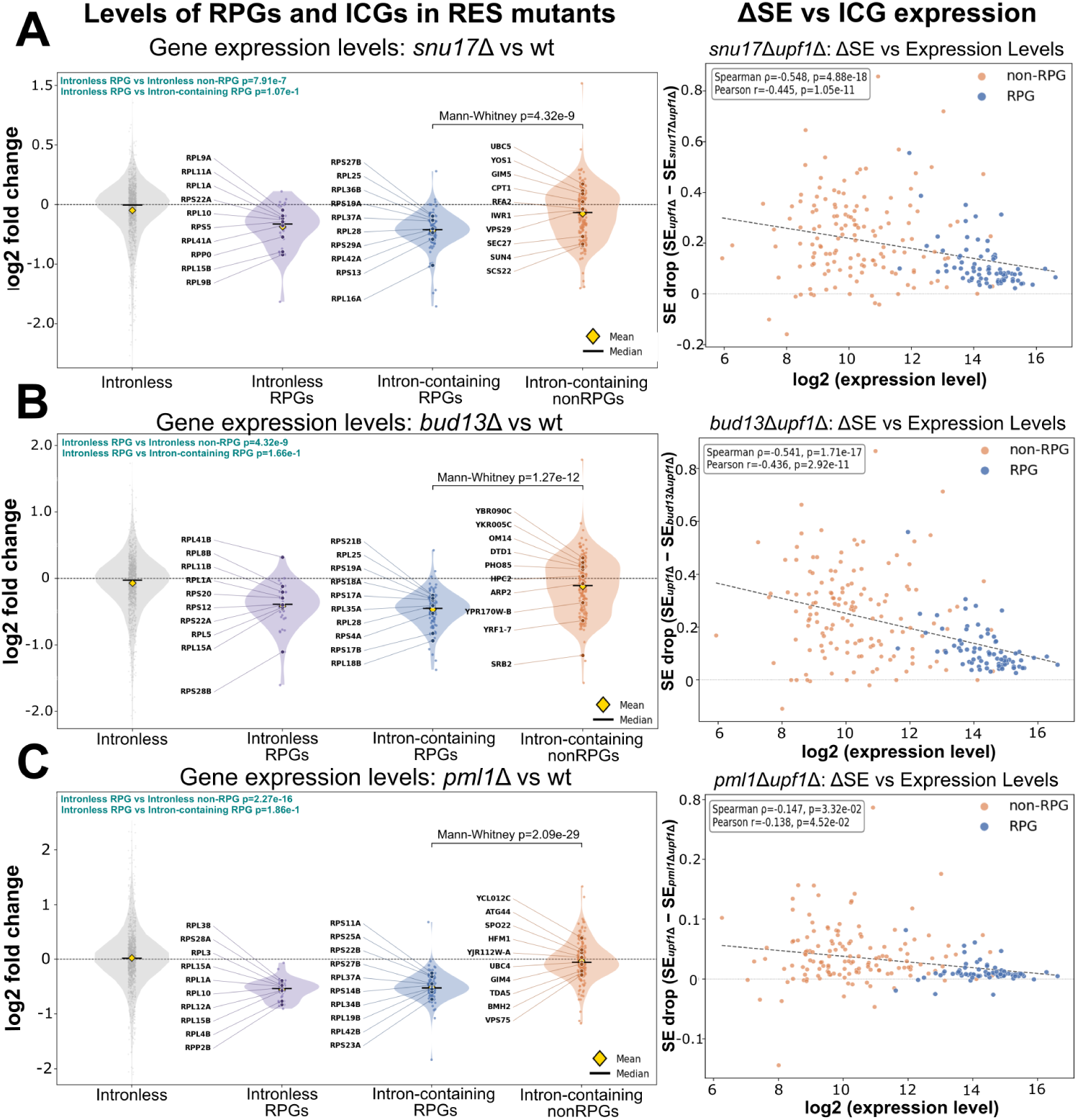
Impact of RES mutants on RNA expression of intron-containing genes. **A.** Left, violin plots of expression change in the *snu17*Δ strain (log2 fold change vs. wt) for intronless genes, intronless RPGs, intron-containing RPGs, and intron-containing non-RPGs. Yellow diamonds and black bars indicate the mean and median of each distribution, respectively. Right: correlation between splicing-efficiency loss and transcript expression in *snu17*Δ*upf1*Δ. Points are colored by gene category (orange, non-RPG; blue, RPG); dashed lines show the linear regression fit for each genotype. **B.** Same as in A for *bud13*Δ and *bud13*Δ*upf1*Δ. **C.** Same as in A for *pml1*Δ and *pml1*Δ*upf1*Δ.

Unexpectedly, *UPF1* deletion alone caused a substantial downregulation of RPGs relative to wt (Supplemental Fig. 3E). Inactivating *SNU17* or *BUD13* in this NMD-deficient background reduces RPG expression only modestly, while *PML1* deletion has no additional effect, raising the possibility that loss of NMD itself engages a feedback mechanism affecting RPG homeostasis in *S. cerevisiae*. This observation is outside the scope of the present study and is not discussed further.

### NMD inactivation rescues the overall transcript levels of several intron-containing genes severely affected by Snu17 or Bud13 deletions

For many of the ICGs downregulated in the RES mutants, their expression was restored in the *upf1*Δ background compared to wt backgrounds (Supplemental Fig. 3F). Some of these likely represent NMD-targets whose unspliced products RES single KOs over-accumulate and are degraded in the cytoplasm by NMD (He et al. 1993). We therefore further analyzed how NMD and the RES complex interact to impact transcript levels of intron-containing genes. To identify the subset of genes for which NMD impacts overall transcripts levels in the absence of each RES subunit, we employed a 2×2 factorial DESeq2 design across our four genotypes (wild type, RESΔ, *upf1*Δ, and RESΔ *upf1*Δ) and extracted the interaction coefficients of each treatment in individual genes: the difference between the RES-deletion effect in the *upf1*Δ background and the RES-deletion effect in the wt background (Love et al. 2014; Muzellec et al. 2023).

Out of the population of downregulated ICGs in *snu17*Δ, *bud13*Δ and *pml1*Δ, a total of 22, 19 and 1 genes, respectively, showed significant RES-NMD interactions (Supplemental Table 3). These NMD-rescued transcripts were predominantly non-RPGs (16 of 22 in *snu17*Δ and 12 of 19 in *bud13*Δ). We curated the list to exclude ICGs with small decreases in SE upon RES inactivation (ΔSE ≤ 0.05). This yielded a set of 17 ICGs in *snu17*Δ and 13 in *bud13*Δ that accumulated mostly as unspliced, whose expression significantly decreased relative to wt, and that harbor PTCs in their unspliced isoforms. Twelve of the 13 *bud13*Δ transcripts were also rescued in *snu17*Δ: *SRB2/MED20*, *HRB1*, *SAR1, EPT1, SEC27*, *TAN1*, *CIN2*, *VPS75*, *ERV1*, *PSP2, RPL16A* and *RPS14B*. *EFB1* was only rescued in bud13Δ while *TDA5*, *YIP3*, *ERV41*, *IST1* and *TFC3* only in *snu17Δ* (Supplemental Fig. 3F). Intron-retention induces a frameshift which introduces a PTC for all these transcripts, with the exception of HRB1 and *RPS14B*, which contain in-frame PTCs within their introns. In the *upf1*Δ background, the decrease of transcript levels detected upon RES subunits KOs was abolished or reversed for 14 out of the 17 transcripts in *snu17*Δ, and all 13 for *bud13*Δ (Supplemental Fig. 3F). In fact, the levels of *HRB1*, *TAN1*, *IST1* and *TFC3* in *snu17*Δ*upf1*Δ and *SRB2* and *HRB1* in *bud13*Δ*upf1*Δ were significantly increased relative to *upf1*Δ, suggesting that the unspliced isoforms of these transcripts accumulate significantly in RES-*upf1*Δ strains. *RPL16A* and *RPS14B,* the only RPGs in this set, were only partially rescued and retained a small decrease upon RES inactivation in both backgrounds. Not surprisingly, no ICGs in *pml1*Δ met this criteria, consistent with the mild splicing phenotype observed relative to the other two strains and potentially compounded by lack of a third replicate. Overall, these results identify a set of mostly non-RPG transcripts whose reduced levels in *snu17*Δ and *bud13*Δ depend on a functional NMD pathway. While this is consistent with their unspliced isoforms being degraded by NMD, the interaction test does not distinguish direct NMD targeting of the unspliced isoform from secondary effects caused by NMD deletion. We thus refer to them as NMD-rescued transcripts rather than *bona-fide* NMD targets.

### Deletion of *PML1* causes growth phenotypes distinct from those of *SNU17* and *BUD13* deletion

Our sequencing data suggests that Pml1p is dispensable for the splicing of nearly all ICGs, whereas Snu17p and Bud13p enhance splicing of many introns. These differences in function should be reflected by differences in growth phenotypes. We therefore followed the growth of all the previously characterized strains in liquid media that includes rich medium (YPD) at 30 °C and 37 °C and in synthetic defined medium (SDC at 30 °C; Fig. 7). From each curve we extracted three parameters: the maximum specific growth rate (µmax), the time to reach OD of 0.5, and the time it took for cells to go from OD 0.25 to 1.5 (the rise time) as a measure of how steeply is cultures moves through the sigmoid (Supplemental Fig. 4; Supplemental table 4). As expected, the *snu17*Δ and *bud13*Δ mutants grew nearly identically under all three conditions. In YPD, their early growth matched that of wt, with a similar µmax and time to OD. 0.5. In SDC, both mutants were impaired from the start, with an µmax 22-24% lower than wt and reaching an OD of 0.5 2 hours later (p < 0.001). *pml1*Δ grew differently from *snu17*Δ and *bud13*Δ in YPD. At 30 °C, *pml1*Δ showed a longer lag phase than the other two mutants (Fig. 7A & C), it had the longest µmax and reached an OD of 0.5 about 1.5 hours after *snu17*Δ and *bud13*Δ. Once growing, however, its rise time matched wt and was about 2 h shorter than that of *snu17*Δ and *bud13*Δ (p < 0.001). As a result, *pml1*Δ overtook both mutants midway through the culture (Fig. 7A), giving a delayed but steep sigmoid rather than the shallow, drawn-out curves of *snu17*Δ and *bud13*Δ. At 37 °C the pattern was stronger: *pml1*Δ was again the last strain to reach OD 0.5, yet it had the shortest rise time of any strain, wt included (p = 0.005). In SDC, *pml1*Δ was indistinguishable from wt for every parameter. The three deletions therefore differ both in the shape of their growth curves and in how that shape depends on the medium.

**Figure 7:**
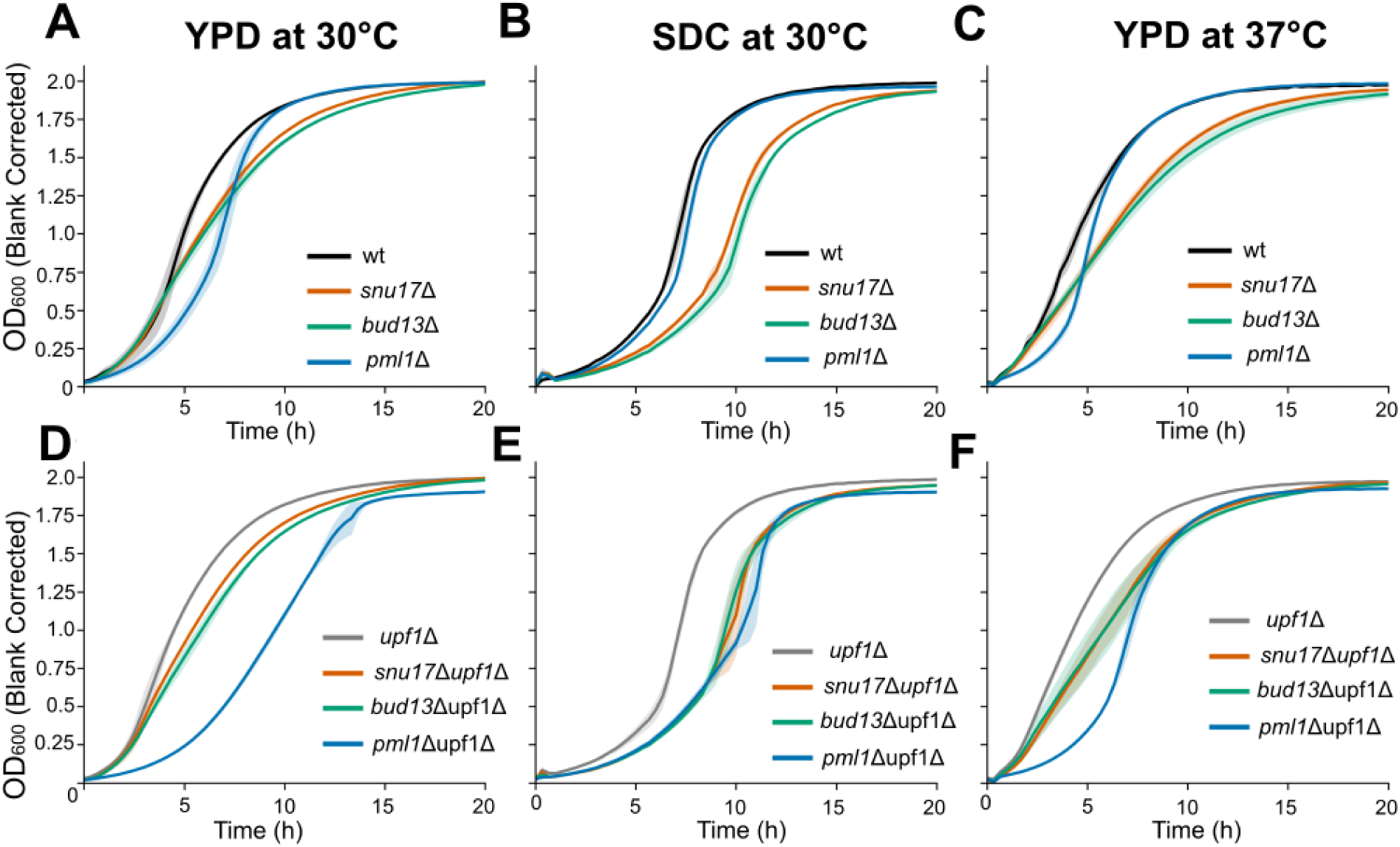
Growth of RES single and RES/*upf1*Δ mutants at 30°C and 37°C. Single RES mutants and wild-type cells grown in A) YPD at 30°C, B) SDC at 30°C and C) YPD at 37°C. Double RES/*upf1*Δ and *upf1*Δ cells grown in D) YPD at 30°C, E) SDC at 30°C and F) YPD at 37°C. Data was collected on the TECAN spark plate reader by measuring OD 600 every 20 minutes for 20 hours with continuous shaking (see materials and methods). Standard deviation is represented by the shadow of each line.

Growth phenotype differences between the mutants in the core complex (Snu17/Bud13) and Pml1 were exacerbated in the NMD-deficient background. *snu17*Δ*upf1*Δ and *bud13*Δ*upf1*Δ behaved like their single mutants: their µmax was close to that of *upf1*Δ, and their rise times were longer. *pml1*Δ*upf1*Δ, in contrast, was markedly sick in YPD. At 30 °C, its µmax was about half that of the other three strains, and it took roughly twice as long to reach OD 0.5 (p ≤ 0.001 for each comparison). At 37 °C, it was again the slowest to reach OD 0.5 (p < 0.001) and had the lowest µmax (p < 0.05), although its rise time stayed close to that of *upf1*Δ. This synthetic sickness depended on the medium. In SDC, all three double mutants grew more slowly than *upf1*Δ, but they did not differ from one another in µmax or time to OD 0.5 (Tukey p ≥ 0.54).

Together, these results show that *snu17*Δ and *bud13*Δ share a growth phenotype that is clearly distinct from that of *pml1*Δ. Deleting *PML1* produces a specific growth signature: a medium-dependent delay followed by a steep transition. It also produces a strong, medium-dependent genetic interaction with *upf1*Δ that we do not observe for the other two subunits. This is notable because *pml1*Δ has the mildest splicing phenotype of the three deletions, which suggests that its growth phenotype may not be caused by global splicing deficiency, but instead due to deficient functions in other processes.

## Discussion

In this study we show that the splicing factors Snu17p and Bud13p work as the core components of the RES complex and that Pml1p acts as an accessory subunit likely serving roles outside of splicing. The results from our functional analysis of the role of these subunits *in vivo* are consistent with prior NMR studies which demonstrated a strong interaction between Snu17p and Bud13p but a weak association of Pml1p to Snu17p (Wysoczanski et al. 2015), and cryo-EM structure of the Bact spliceosome, which showed that yeast Snu17p binds directly the pre-mRNA, forming a heterodimer with Bud13p that sits within the main body of the spliceosome (Yan et al. 2016; Fig. 1A & D). Pml1p, on the other hand, sits mainly outside the main body of the spliceosome (Wysoczanski et al. 2015; Fig. 1C & D).

### Pml1 is not required for efficient splicing or unspliced pre-mRNA retention genome-wide

In our study, Pml1p deletion results in phenotypes clearly distinct from those observed upon deletion of Snu17p and Bud13p. Genome-wide, the SEs of annotated introns remain largely unaffected in *pml1*Δ*upf1*Δ (Fig. 1A) while deletion of Snu17p and Bud13p appear to be phenotypically identical as shown by PCA analysis and result in widespread increase in intron retention, particularly in non-RPGS (Fig. 1B). The growth phenotypes reinforce the conclusion that Pml1p differs functionally from Snu17p and Bud13p. The *snu17*Δ and *bud13*Δ mutants grow almost identically, consistent with their acting as a single unit within the spliceosome while the *pml1*Δ strain instead shows a delayed but steep growth curve in rich medium and is indistinguishable from wt in SDC. The observation that its growth is delayed more than *snu17*Δ or *bud13*Δ in YPD, despite having the mildest splicing defect, suggests to us that the *pml1*Δ phenotype is not a consequence of general splicing deficiency. The observed synthetic sickness in *pml1*Δ*upf1*Δ is likely not caused by functional redundancy between Pml1p and Upf1p, as we did not detect any increase in known Upf1p targets in *pml1*Δ (Supplemental table 2). The leakage of unspliced pre-mRNAs previously described in the *pml1*Δ strain (Dziembowski et al. 2004) suggested that Pml1 and Upf1 could work in parallel pathways involved in the detection and degradation of unspliced or improperly processed species. However this result was obtained with a reporter containing an artificial intron, and whether or not Pml1 acts to retain endogenous unspliced transcripts was not investigated. The lack of increase of unspliced pre-mRNAs in the *pml1*Δ*upf1*Δ strain compared to the single *upf1*Δ mutant is inconsistent with a global role for Pml1 in retaining unspliced precursors in the nucleus.

### Snu17 and Bud13 form the functional core of the RES complex required to enhance splicing of weak introns

Our data does conclusively demonstrate, on the other hand, that Snu17p and Bud13p are critical to maintaining splicing efficiency of introns that are inefficiently spliced at background levels. *In vitro* studies using *snu17*Δ and *bud13*Δ extracts demonstrate that the spliceosome can carry out both transesterification reactions in the absence of either one of these two RES complex components, albeit in a structurally compromised state (Bao et al. 2017). Non-RPGs with short introns that lack consensus 5′ss and bp sequences can’t be efficiently spliced as the spliceosome undergoes large structural rearrangements from the A complex to the P complex in the absence of Snu17p and Bud13p. Deviations of branch-point sequences from the consensus sequence in RPGs, however, cannot fully explain splicing defects, as the first introns in *RPL7A* and *RPL7B* lack a consensus bp sequence and yet they are among the most efficiently spliced introns in *snu17*Δ*upf1*Δ and *bud13*Δ*upf1*Δ (Supplemental Table 1; Supplemental Fig. 3D). This mirrors our finding that RPGs with non-consensus 5′ss are spliced as efficiently as those with a consensus 5′ss (Supplemental Fig. 2B), which indicates that RPG resistance by RES loss is not conferred by the sequence of the splice signals themselves. Instead, the significantly larger introns and the larger TSS-to-3′ss distances that differentiate RPGs from non-RPGs (Supplemental Fig. 1A) are the more likely determinants of this resistance, consistent with their ranking as strong predictors of high SE in our Random Forest Analysis (Fig. 3A, B & C).

### A role for polyU tracts in enhancing splicing efficiency prior to the second chemical step of splicing in S.cerevisiae

In mammalian introns, the polypyrimidine tract plays a critical role in early intron recognition through binding of U2AF (Sickmier et al. 2006). In *S.cerevisiae*, polypyrimidine sequences present between the branchpoint and the 3′ss have been shown to impact 3′ss selection (Patterson & Guthrie 1991). The data we provide show that poly(U) tracts present between the branchpoint and the 3′ss globally contribute to splicing efficiency, as their presence buffers splicing efficiency against the loss of Snu17p or Bud13p. This result is consistent with data obtained from a library of synthetic introns showing that the presence of U-tracts is associated with higher splicing efficiencies (Schirman et al. 2021). Because of the overall binding pattern of Snu17p reanalyzed from crosslinking data (Fig. 1E), we propose that these sequences may play a role in promoting efficient spliceosome assembly, either at an early stage of intron recognition, or by helping maintain the RES complex at the vicinity of the pre-mRNA. To our knowledge, these data provide for the first time evidence that poly(U) tracts may play a role in splicing at a stage earlier than the 2nd chemical step of splicing.

### Global impact of the RES complex on cellular fitness

Beyond its role in splicing, our data reveal that loss of either RES-complex subunit triggers a broad transcriptional downregulation of RPGs independent of their splicing status: intronless RPGs are repressed to the same extent as intron-containing ones (Fig. 6A, B & C). This points to a feedback mechanism that reduces RPG expression genome-wide. We don’t believe this effect is unique to deletion of RES complex components but rather is a general stress response to spliceosomal perturbations (Shore et al. 2021). This is supported by the observation that *PML1* deletion leads similarly to a decrease of intron-containing and intronless RPGs levels despite having little effect on global SEs (Fig 1A; Fig 6A).

Together, our results support a model in which Snu17p and Bud13p act as a structural unit that buffers the spliceosome against intrinsically weak splicing signals—short introns, non-consensus bp sequences, and poorly defined polyU tracts—without being strictly required for catalysis itself. This is consistent with the observed residual splicing activity of *snu17*Δ and *bud13*Δ extracts *in vitro* (Bao et al. 2017). Pml1p, by contrast, is largely dispensable for this buffering function and potentially contributes to fitness through a distinct pathway that appears to be NMD dependent as the *pml1*Δ*upf1*Δ grows remarkably sicker than the other two double mutant strains (Fig. 8D open question that remains is what roles exactly is Pml1p serving outside of splicing and what process underlies the observed synthetic sickness of *pml1*Δ*upf1*Δ even in the absence of any major splicing defects.

## Materials and Methods

### Yeast strains and strain construction

The wild-type strain was BY4742 (*MATα his3Δ1 leu2Δ0 lys2Δ0 ura3Δ0*). The *snu17*Δ*::kanMX4*, *bud13*Δ*::kanMX4* and *pml1*Δ*::kanMX4* single mutants in this background came from the Open Biosystems yeast deletion collection. To make the RES/*upf1Δ* double mutants, we PCR-amplified the *upf1Δ::HIS3* cassette from an existing strain along with its flanking *UPF1* sequence, from genomic DNA of this strain (primers: 5′-GTCACTAGCGAGTTCATTCTCCAATTG-3′ and 5′**-**CGAGAGAGAAAAGAGGAGTTCTAAG-3′). We then integrated it at the *UPF1* locus of each single mutant by lithium acetate transformation (Gietz and Schiestl 2007). Transformants were selected on synthetic complete medium lacking histidine. Correct integration was confirmed by diagnostic PCR (primers: 5′-GGAAGGGCAGCAAGACCG-3′ and 5′**-**GCAGCGCTCATTTCACGG-3′).

### Growth Assays and analysis

Overnights of wt, *upf1*Δ, RES single and RES-NMD double KOs were grown for 18 hours to saturation. Cells were backdiluted in 50 mL of pre-warmed media to an O.D. of 0.1 and allowed to grow to two doublings (O.D. of 0.4). Cell cultures were then back-diluted to a final O.D of 0.05 in a total value of 200 uL of the corresponding stress media in a 96-well plate. Cell growths and data acquisition were performed using the TECAN Spark Multimode microplate reader and its corresponding TECAN SparkControl software. Briefly, the protocol consisted of prewarming (or cooling) the apparatus half an hour before the first data point acquisition. The kinetic loop was set for at least 81 cycles with a cycle representing data point measurements performed every 20 minutes. The measurement wavelength was set to 600 nm with a bandwidth of 3.5. The shaking was done continuously, in orbital mode, with an amplitude of 2.5 mm and a frequency of 216 rpms. OD readings were blank-corrected and maximum specific growth rate (µmax, h⁻¹) of each replicate was estimated as the steepest slope of ln(OD₆₀₀) versus time. Slopes were obtained by linear regression over all windows of six consecutive time points (100 min) lying entirely within OD₆₀₀ 0.1–1.0. As a way to describe curves shapes, we calculated the maximum specific growth rate (µmax) as the steepest slope of ln(OD₆₀₀) versus time, from linear regression over all six-point windows (100 min) lying entirely within OD₆₀₀ 0.1–1.0, adapted from Hall et al. (2013). We also calculated the time it took cultures to reach an OD of 0.5 and the time it took cultures to increase from an OD of 0.25 to 1.5 (rise time); these two points were chosen since they are both within the exponential growth phase and before the deceleration phase (Hall et al. 2013). Parameters were calculated per replicate (n = 3) and are reported as mean ± SD. Within each growth condition, genotypes were compared within each background (wt and single deletions; *upf1*Δ and double deletions) by one-way ANOVA followed by Tukey’s HSD.

### RNA extractions and library preparations

Overnights of all strains were grown in replicates at 30 °C in YPD (1% w/v yeast extract, 2% w/v peptone, and 2% w/v dextrose, at 30°C) for 18 hours. Overnights were backdiluted the next day to an O.D of 0.1 and allowed to grow to two doublings to an O.D. ∼0.45. Cells were harvested by centrifuging at 3000 RPMs for 4 minutes at 1800 g, flash frozen in liquid nitrogen and stored at 80 30 °C. Total RNAs were extracted using the hot phenol chloroform method and treated with DNase I (NEB; M0303S). 1 ug of total DNase treated RNA for each replicate was ribodepleted with RiboCop rRNA Depletion Kit for yeast (Lexogen; 190) and the total output was used for NGS library preparation using the CORALL RNA-seq V2 library prep kit (Set B, UDI12B_0001-0096; Lexogen, Vienna, Austria) following the manufacturer’s protocol. Libraries were then sequenced using the Illumina NovaSeq 6000 in paired-end mode.

### RNA sequencing and Bioinformatics analysis

RNA sequencing libraries were prepared with the CORALL Total RNA-Seq Library Prep Kit V2 (Lexogen) and sequenced paired-end. Each Read 1 begins with a 12-nt unique molecular identifier (UMI), which was removed from the read and appended to the read name with umi_tools extract (v[1.1.6]; Smith et al., 2017). Adapter sequences were then removed from Read 1 (AGATCGGAAGAGCACACGTCTGAACTCCAGTCAC) and Read 2 (AGATCGGAAGAGCGTCGTGTAGGGAAAGAGTGT) with cutadapt (v[5.2]; Martin, 2011). Bases with quality below 20 were trimmed from the 3′ ends. Read pairs in which either read was shorter than 20 nt after trimming were discarded.

### Sequences Alignment

Trimmed, UMI-labeled read pairs were aligned to the *S. cerevisiae* S288C reference genome (assembly R64-5-1) using BBMap. A maximum indel size of 1200 nt (maxindel=1200) and a maximum insert size of 1200 nt (pairlen=1200) were specified to permit gapped alignments spanning annotated and unannotated introns; gaps of 20 nucleotides or greater were reported as intron-spanning (N) segments in the resulting CIGAR strings (intronlen=20). Resulting alignments were coordinate-sorted and indexed with samtools.

PCR duplicates were then removed from each sample’s sorted BAM file with umi_tools dedup (Smith et al., 2017) in paired-end mode (--paired), which collapses read pairs sharing identical mapping coordinates and identical (or network-adjacent, under the default directional deduplication method) UMIs into a single representative pair. Deduplicated alignments were re-sorted and indexed with samtools prior to downstream analysis.

### DEseq2 analysis

Gene-level read counts were generated with featureCounts (Liao et al. 2014) and analyzed with PyDESeq2 (Muzellec et al. 2023), a Python implementation of DESeq2 (Love et al. 2014). Genes with fewer than 10 reads summed across all samples were removed. For genome-wide comparisons, all eight genotypes were fitted in a single model (∼ genotype), so dispersions were estimated from the full dataset. Each RES single mutant and *upf1Δ* was compared with wt, and each RES/*upf1*Δ double mutant with *upf1*Δ. Significance was assessed by Wald test with Benjamini–Hochberg correction (adjusted p < 0.05). Log2 fold changes are reported without shrinkage. We tested whether ICGs were preferentially downregulated by comparing the fraction of significantly downregulated genes among ICGs and intronless genes with Fisher’s exact test. RPGs were identified from the SGD annotation (R64-5-1) and split into intron-containing (n = 90) and intronless (n = 48) genes. Their log2 fold changes were compared by [Mann–Whitney U test].

### Random Forest Permutation

Intronic features predictive of splicing efficiency were identified by Random Forest regression (scikit-learn v[1.7.2]; 500 trees, minimum of five introns per leaf). A separate model was fit for each genotype, using mean SE across replicates as the response. Ten predictors were computed for each intron: intron length; the number of mismatches to the consensus 5′ss (GUAUGU) and branchpoint (UACUAAC) sequences; the distance from the branchpoint adenosine to the 3′ss; the uridine fraction and longest uridine tract between the branchpoint and the 3′ss; the distances from the dominant TSS (McMillan et al. 2019), to the 5′ss, branchpoint and 3′ss; and ribosomal protein gene status. Branchpoints were curated manually and validated against the Branch-seq map of Gould et al. (2016). Of the 200 introns detected by Branch-seq, 193 matched a mapped branchpoint. The remaining seven carry a consensus UACUAAC that Branch-seq did not detect and were retained. Introns without Branch-seq data were assigned by best match to UACUAAC near the 3′ss. Introns lacking any predictor were excluded.

Model performance was evaluated by 5-fold cross-validated R², and significance was assessed against a null distribution from 100 permutations of the response. Feature importance was measured as the mean decrease in R² after 50 independent permutations of each feature, and its significance was tested with a one-sided Wilcoxon signed-rank test with Benjamini–Hochberg correction. The direction of each feature’s effect was assigned from its Spearman correlation with SE. Because the three TSS distances are correlated, their importances should be considered together.

### Sequence logos

Introns were classified for each RES double mutant relative to *upf1*Δ, among those with SE ≥ 0.30 in *upf1*Δ. Introns with |ΔSE| < 0.10 were classified as not affected. Introns with ΔSE ≥ 0.30 and a significant change between strains (Welch’s t-test, p < 0.05) were classified as affected. All others were classified as intermediate. Sequences of introns were extracted from the S288C reference genome (R64-5-1), using branchpoints defined as described above. Logos were generated for the 5′ss (first 6 nucleotides of the intron), the branchpoint (from position -6 to position +2 relatrive to bp A) and the 3′ss (the 10 nt upstream of the terminal AG, including AG). A reference set of all canonical introns quantified in every strain was included for comparison. Logos show information content (bits) and were plotted with Logomaker (Tareen & Kinney; 2020)

### RES × NMD interaction analysis

To identify transcripts whose downregulation in RES mutants depends on NMD, we fitted a separate model for each RES subunit. Each model used the four relevant genotypes (wt, RES single deletion, *upf1*Δ, and the RES/*upf1*Δ double deletion) and a 2 × 2 factorial design (∼ RES + NMD + RES:NMD), with wt as the reference for both factors. In this model, the RES:NMD interaction coefficient equals the RES-deletion effect in the *upf1*Δ background minus its effect in the wt background. We tested it directly by Wald test. NMD-target candidates had to meet three criteria: significant downregulation in the RES single mutant relative to wt (adjusted p < 0.05); a significant positive interaction (adjusted p < 0.05), meaning the downregulation is attenuated when NMD is inactivated; a reduction in mean splicing efficiency of more than 0.05 in the RES single mutant relative to wt. The ICG expression categories in Figure 2D (up, down, unchanged) were taken from the RES-deletion coefficient of these same models. For each high-confidence candidate, we checked for premature termination codons by reconstructing the intron-retained coding sequence from the R64-5-1 genome annotation.

## Data Availability

Raw RNA sequencing data for all strains and processed splicing efficiency and gene count matrixes have been submitted to the NCBI Gene Expression Omnibus GEO (https://www.ncbi.nlm.nih.gov/geo/) under accession number GSE350189. All custom scripts used in this study can be found at (https://github.com/AArce-Lab/RES_project).

## Acknowledgements

We thank everyone in the Chanfreau Lab for the discussions and Isaac Yang for comments on the manuscript. We would also like to thank Martina Rivas for her assistance in preparing culture media and maintaining laboratory supplies, which streamlined our experimental workflows.

Supported by grant GM130370 from NIGMS (GFC) and by the National Institute of General Medical Sciences of the National Institutes of Health under Award Number T32GM145388.

